# Shedding light on human olfaction: electrophysiological recordings from sensory neurons in acute slices of olfactory epithelium

**DOI:** 10.1101/2023.03.27.534398

**Authors:** Andres Hernandez-Clavijo, Cesar Adolfo Sánchez Triviño, Giorgia Guarneri, Chiara Ricci, Fabian A. Mantilla-Esparza, Kevin Y. Gonzalez-Velandia, Paolo Boscolo-Rizzo, Margherita Tofanelli, Pierluigi Bonini, Michele Dibattista, Giancarlo Tirelli, Anna Menini

## Abstract

The COVID-19 pandemic brought attention to our limited understanding of human olfactory physiology. While the cellular composition of the human olfactory epithelium is similar to that of other vertebrates, its functional properties are largely unknown. We prepared acute slices of human olfactory epithelium from nasal biopsies and used the whole-cell patch-clamp technique to record electrical properties of cells. We measured voltage-gated currents in human olfactory sensory neurons and supporting cells, and action potentials in neurons. Additionally, inward currents and action potentials responses of neurons to a phosphodiesterase inhibitor indicated that the transduction cascade involves cAMP as a second messenger. Furthermore, responses to odorant mixtures demonstrated that the transduction cascade was intact in this preparation. This study provides the first electrophysiological characterization of olfactory sensory neurons in acute slices of the human olfactory epithelium, paving the way for future research to expand our knowledge of human olfactory physiology.

## Introduction

Human olfaction has long been considered a neglected sense and the recent COVID-19 pandemic has highlighted the scarce knowledge we have of human olfactory physiology. The sudden and widespread olfactory loss experienced during the pandemic caught us unprepared, with many individuals struggling to recover their sense of smell^1–7^. Indeed, although the morphology of the human olfactory epithelium was well known, there was limited knowledge about the molecular and functional landscape of different cell types within the human olfactory epithelium. Molecular data became soon available in the early phase of the pandemic^8–11^ but the functional properties of the cells are still largely unknown.

The human olfactory epithelium is located in the upper posterior part of the nasal cavity, and it is found in patchy regions that alternate with non-sensory epithelium. Its cellular composition and organization are similar to that of most other vertebrates^12–15^. The epithelium consists of three main cell types: olfactory sensory neurons, supporting (or sustentacular) cells, and basal cells. Olfactory sensory neurons are bipolar neurons that have one dendrite ending with a knob from which several cilia originate at the surface of the epithelium, a soma and a single axon reaching the olfactory bulb. Supporting cells, the “unsung heroes”^10^ of the olfactory epithelium, are columnar in shape, extending from the basal to the apical portion of the epithelium. They provide structural support to olfactory sensory neurons, and bear microvilli on their apical side^16^. Basal cells are located at the basal part of the epithelium and have the ability to regenerate various cell types within the olfactory epithelium^11,16,17^.

Recent research during the COVID-19 pandemic has identified supporting cells as the primary target of SARS-CoV-2 in the olfactory epithelium^8–10^. Furthermore, the compromised functionality of supporting cells, along with inflammatory processes, can exacerbate olfactory loss^18^.

Olfactory transduction in rodents and amphibians has been extensively studied and it is well-established that it occurs in the cilia of olfactory sensory neurons^19–23^. This process begins with the binding of odorant molecules to specific G protein-coupled odorant receptors, which activate a biochemical cascade that increases cAMP concentration within the cilia. As a result, the open probability of cyclic nucleotide-gated (CNG) channels increases, allowing the entry of Na^+^ and Ca^2+^ and inducing neuron depolarization^24–27^. The increase in Ca^2+^ concentration within the cilia then activates the Ca^2+^-activated Cl^-^ channel TMEM16B (also named ANO2) that contributes to regulate neuron depolarization^28–34^. When the depolarization reaches the threshold, action potentials are generated and transmitted to the olfactory bulb^35–37^. In combination with calmodulin, Ca^2+^ also contributes to response termination by enhancing the activity of the phosphodiesterase PDE1C2, which hydrolyzes cAMP, reducing the open probability of CNG channels^38,39^. The use of the PDE inhibitor 3-isobutyl-1-methylxanthine (IBMX) has unveiled the presence of a basal cAMP concentration, as upon IBMX application inward currents were measured in the whole-cell voltage-clamp configuration in olfactory sensory neurons in amphibians and rodents^40–42^. Basal cAMP fluctuations (hence the IBMX response) are caused by the constitutive activity of odorant receptors that activate the transduction cascade, producing a depolarization followed by action potential generation. Different odorant receptors show different levels of constitutive activity^43–45^.

The mature olfactory sensory neurons, which express the olfactory marker protein (OMP)^46,47^ and only one odorant receptor type among about 400-1000^10,48^, are the main functional units in the olfactory epithelium in most vertebrates, including humans. In human mature olfactory sensory neurons, some genes coding for proteins known to be involved in odorant signal transduction in rodents are expressed, such as several odorant receptor genes, G protein alpha (*GNAL*) and gamma subunits (*GNG13*), adenylyl cyclase type 3 (*ADCY3*), cyclic nucleotide gated channel alpha2 (*CNGA2*), and the calcium-activated chloride channel (*TMEM16B*/ANO2)^8–11,49^. However, while immunohistochemistry data have confirmed expression of the G protein alpha and gamma subunits in human olfactory sensory neurons^11,13^, no data have been published for the other proteins potentially involved in the human olfactory transduction cascade.

From a functional point of view, a pioneering study by Restrepo et al.^50^ reported that viable human olfactory sensory neurons could be dissociated from olfactory tissue biopsies, and showed, by using Ca^2+^ imaging, that some neurons responded to odorants with an increase in intracellular Ca^2+^ concentration. Further studies by the same laboratory^51,52^ showed that some human olfactory neurons also responded to odorants with a decrease in intracellular Ca^2+^ concentration, a response never observed in neurons from other vertebrates^50,53–55^, suggesting that human olfactory neurons have unique properties compared to other vertebrates. Moreover, Gomez et al.^56^ found that protein kinases A and C modulate odorant responses in different ways in human and rat olfactory neurons, indicating additional differences between the two species. Overall, these studies provided insight into the odorant-induced Ca^2+^ changes in human olfactory neurons revealing differences from other vertebrates.

In rodents and amphibians, electrophysiological techniques have been extensively used to study the functional properties of olfactory sensory neurons while only a few studies have been reported in humans^50,57,58^. One of these studies used the inside-out patch-clamp technique from the dendritic knob of dissociated human olfactory neurons to characterize activation of CNG channels by cAMP, providing evidence that these channels may be involved in olfactory transduction in humans^57^. Two other reports investigated the electrical properties of isolated human olfactory neurons with the whole-cell patch-clamp technique^50,58^. Both studies consistently measured outward voltage-gated currents in response to depolarizing voltage steps, while transient inward currents were rarely observed in human olfactory neurons. The absence of transient inward currents in most human olfactory neurons is surprising, as they are found in other vertebrates^59^, and this may be due either to a unique aspect of human olfactory transduction^58^ or to neuron damage during the dissociation procedure^50^.

Knowledge of the initial electrical events in human olfactory neurons is crucial for understanding the signals transmitted from the periphery to the brain. To achieve this, it is essential to use a preparation that closely mimics the physiological environment of human olfactory sensory neurons. In our study, we developed a technique to obtain acute slices from biopsies of the human olfactory epithelium, which provides a more physiological setting for olfactory neurons than dissociated cells. Using this preparation, we employed the whole-cell patch clamp technique to measure the basic biophysical properties and voltage-gated currents of olfactory sensory neurons and supporting cells, and recorded action potential generation in olfactory neurons. Moreover, we were able to record IBMX and odorant-induced transduction currents, providing the first functional characterization of olfactory sensory neurons from acute slices of the human olfactory epithelium.

## Results

### Immunohistochemistry of the human olfactory epithelium

We performed an immunohistochemical analysis of biopsies of human nasal tissues using specific markers to identify the presence of olfactory sensory neurons and supporting cells.

We used β-tubulin III (TUJ1) a known marker for neurons and the olfactory marker protein (OMP) to stain mature olfactory sensory neurons and clearly identified the typical bipolar morphology of olfactory sensory neurons (Fig. 1a) with cell bodies within the epithelium and a dendrite extending till the apical side. Axon bundles were also distinguishable under the basal lamina. To identify supporting cells, we used ERMN as their marker^13,15^ and observed a staining of the apical part of the epithelium, mutually exclusive with TUJ1 staining (Fig. 1d).

**Fig. 1.**
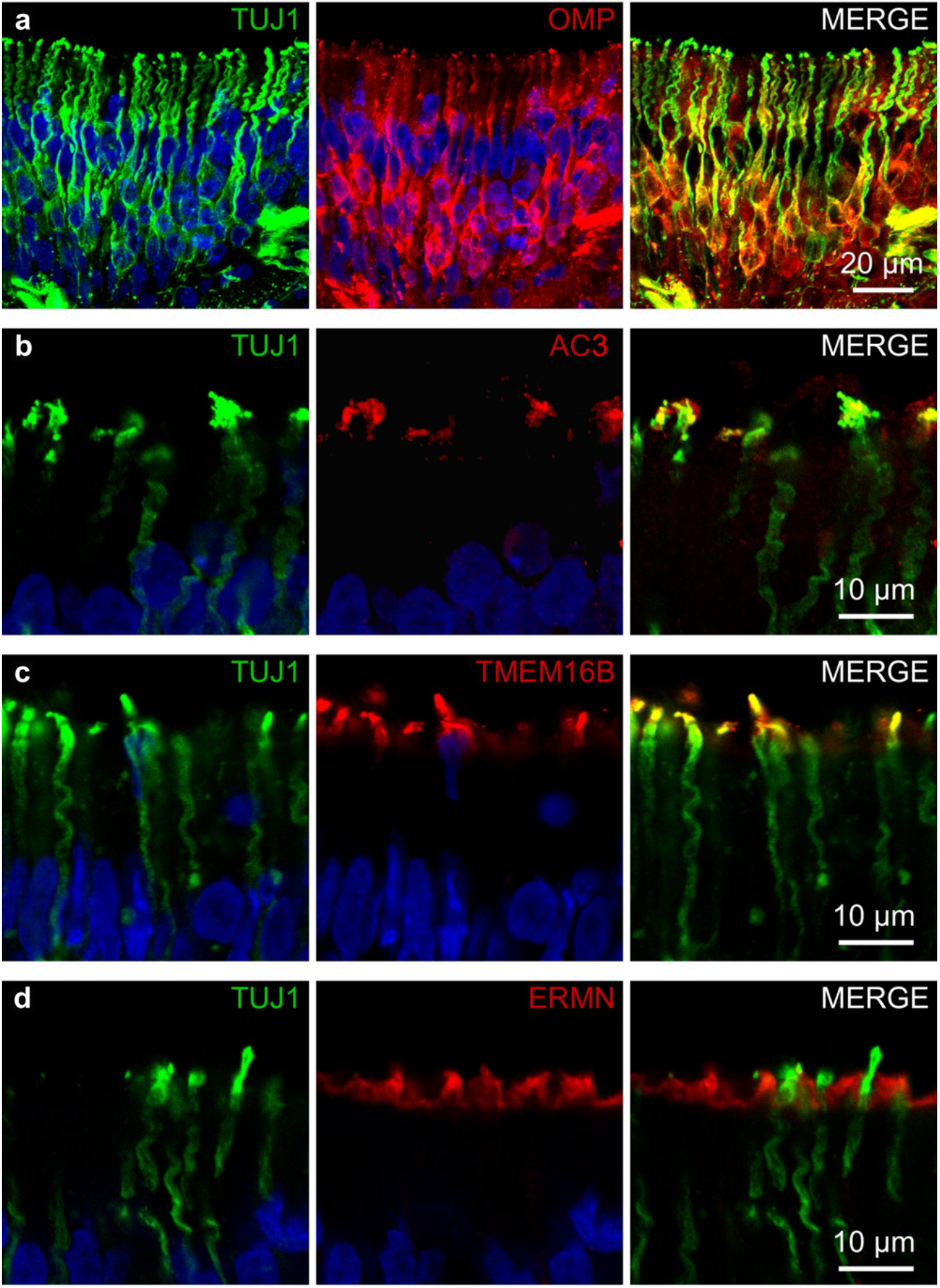
Olfactory sensory neurons from human olfactory epithelium express signal transduction proteins at the apical part. **a** Olfactory sensory neurons stained with the neuronal marker TUJ1 (green) and OMP (red). **b** Co-expression of TUJ1 (green) and AC3(red) at the apical part of olfactory sensory neurons. **c** Co-expression of TUJ1 (green) and TMEM16B (red) at the apical part of olfactory sensory neurons. Note that AC3 and TMEM16B staining are present only in the dendritic knob and ciliary region of the neuron. **d** Non-overlapping staining for the neuronal marker TUJ1 (green) and ERMN (red), a marker for the apical region of supporting cells. Cell nuclei were stained with DAPI (blue).

Although it is well known that in rodents several proteins of the transduction cascade are expressed in the apical dendritic knob and ciliary regions of olfactory sensory neurons, in humans the expression and cellular localization of most of these proteins have not been investigated yet. Immunostaining with antibodies against adenylyl cyclase type 3 (AC3) and the Ca^2+^-activated Cl^-^ channel TMEM16B revealed the expression of both proteins in the apical knob and ciliary region of the TUJ1 positive neurons (Fig. 1b,c).

These results extend previous immunohistochemistry data showing that AC3 and TMEM16B are localized in the dendritic knob and cilia of human olfactory sensory neurons, where olfactory transduction takes place.

### Voltage-gated currents in human olfactory sensory neurons and supporting cells

Since a very limited number of studies have attempted to measure the electrophysiological properties of human olfactory sensory neurons and these studies have been performed only on isolated neurons, we asked whether it is possible to record the electrical activity from cells in acute slices of the human olfactory epithelium. Slices may provide a more physiological environment to the olfactory neurons and better cell viability, important for obtaining long lasting and stable recordings.

We first established the viability of obtaining electrophysiological recordings by measuring basic electrical properties and voltage-gated currents in the whole-cell voltage-clamp configuration. To visually identify cells, we dissolved Alexa Fluor 594 in the intracellular solution filling the patch pipette and took fluorescence images after obtaining the whole-cell configuration and the diffusion of the fluorophore inside the cell (Fig. 2 a,d). A human olfactory sensory neuron, with its typical morphology comprising a cell body toward the basal part and a dendrite extending to the apical part of the epithelium, is shown in Fig. 2a, demonstrating that it is possible to reach a whole-cell configuration and to visually identify neurons in acute slices of the human olfactory epithelium. We then evaluated the resting membrane potential in current-clamp at I = 0 in neurons and calculated an average value of -52 ± 5 mV (range -76 to -24 mV, n = 13). The membrane input resistance, estimated in voltage-clamp, had an average value of 4.2 ± 1.2 GΩ (range 1.0 to 9.7 GΩ, n = 12).

**Fig. 2.**
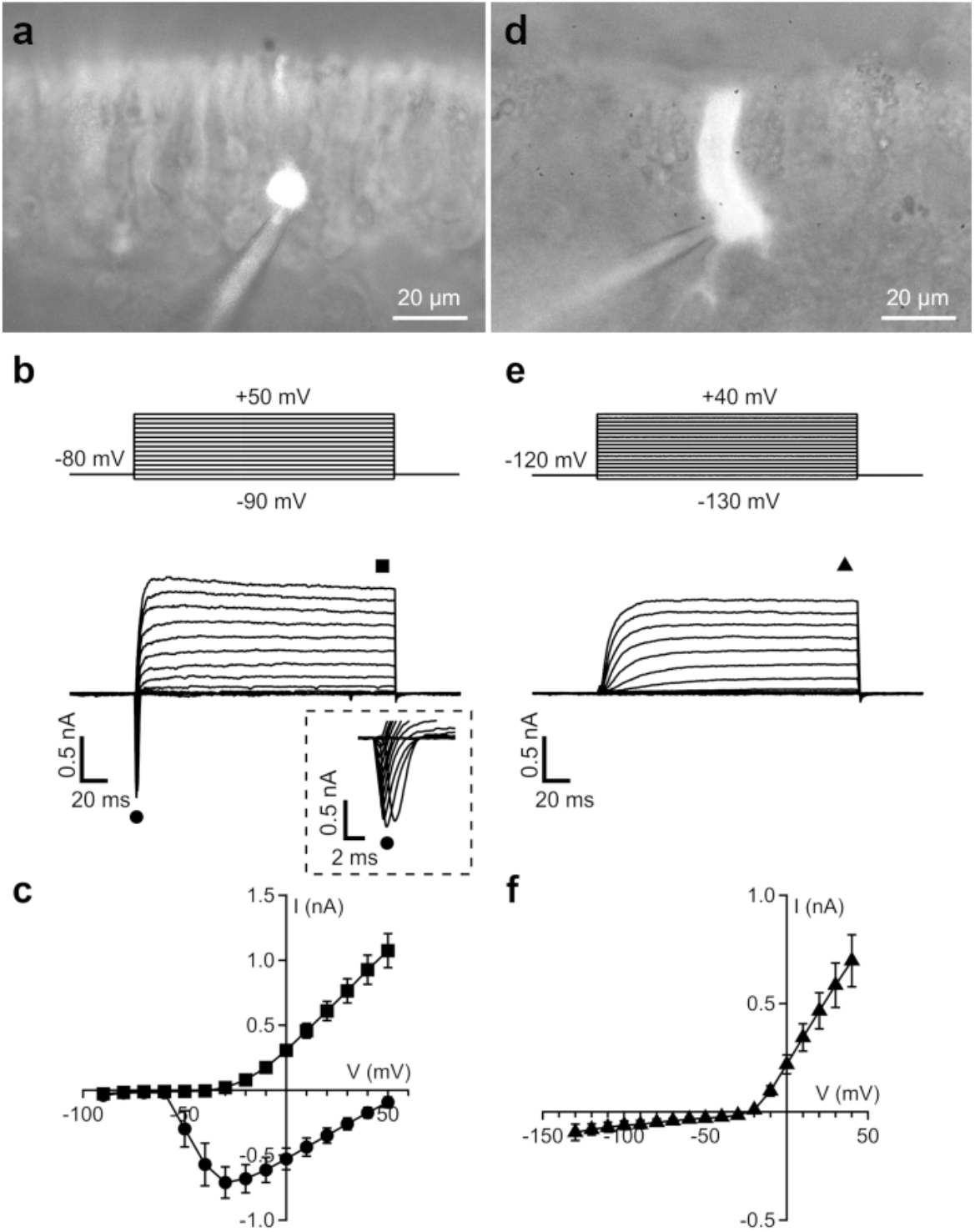
Voltage-gated currents in olfactory sensory neurons and supporting cells from acute slices of the human olfactory epithelium. **a**,**d** Fluorescence micrographs of an olfactory sensory neuron (**a**) and a supporting cell (**d**) filled with Alexa Fluor 594 through the patch pipette. **b**,**e** Representative whole-cell currents recorded using the voltage protocols indicated at the top of the panels. The holding potential was −80 mV for olfactory sensory neurons (**b**) and -120 mV for supporting cells (**e**). Voltage steps in 10 mV increments were applied. The inset in (**b**) shows details of the inward currents on an expanded time scale. **c**,**f** Plot of average ± SEM amplitudes of inward (black circles) and outward (black squares) currents in olfactory sensory neurons (**c**, n = 10) and outward (black triangles) currents in supporting cells (**f**, n = 12) versus the test potential.

Next, we measured voltage-gated currents in human olfactory sensory neurons. Transient inward currents followed by outward currents were activated upon depolarization from a holding potential of –80 mV (Fig. 2b). Current–voltage relations were measured at the peak of the inward currents or at the end of the sustained outward currents, averaged from several neurons and plotted in Fig. 2c. The average current-voltage relations show that the transient inward current activated between –60 and –50 mV and reached a peak at -30 mV, with an average value of -0.7 ± 0.1 nA (n = 10) and then decreased toward 0 between 50 and 60 mV. Outward currents activated at about -30 mV and increased their amplitude with the depolarizing step potential reaching an average value of 1.1 ± 0.1 nA (n = 10) at +50 mV (Fig 2c).

We also recorded from supporting cells in the whole-cell voltage-clamp configuration. Fluorescence images of supporting cells showed their typical columnar shape with fine processes extending toward the basal part of the epithelium (Fig. 2d). The average membrane input resistance was 2.2 ± 0.6 GΩ (range 0.5 to 7.1 GΩ, n = 12). In a first set of experiments, we elicited voltage-gated currents with the same step protocol used for neurons from a holding potential of -80 mV and measured sustained outward currents (data not shown). As we and others have previously shown that supporting cells in mice also have voltage-activated transient inward currents^60,61^, in a second set of experiments we lowered the holding potential to -120 mV before applying a depolarizing step protocol, also in this condition no transient inward currents were measured (Fig. 2e). Outward currents activated at about -10 mV and increased with the depolarizing step potential (Fig. 2f) reaching an average value of 0.7 ± 0.1 nA (n = 12) at +40 mV.

These electrophysiological data show that voltage-gated currents in human olfactory sensory neurons have both transient inward currents and outward currents as in other vertebrate species. On the other side, human supporting cells displayed only outward voltage-gated currents, differently from mice, where also transient inward currents have been reported.

### Firing patterns of human olfactory sensory neurons

To investigate the firing patterns of human olfactory sensory neurons in the acute slice preparation we used whole-cell current-clamp recordings. The responses of three representative neurons to current injections from -2 to 10 pA of 2 s duration show the different types of spiking patterns we measured (Fig. 3a-c). The first neuron in Fig 3a generated a tonic firing consisting of sustained train of action potentials in response to current injection of 2 pA and displayed an increasing number of spikes up to 6 pA current steps. At higher current injections of 8 and 10 pA, the same neuron generated a phasic firing with a brief train of action potential of decreasing amplitude followed by small oscillations around a voltage plateau (Fig. 3a). The other two neurons fired only one or two action potentials in response to current injections of 2-4 pA up to 10 pA, followed by a voltage plateau (Figs. 3b,c). Of eight neurons, two displayed tonic firing at current steps between 2 and 6 pA followed by phasic firing at 8 and 10 pA, while six fired only one or a few action potentials.

**Fig. 3.**
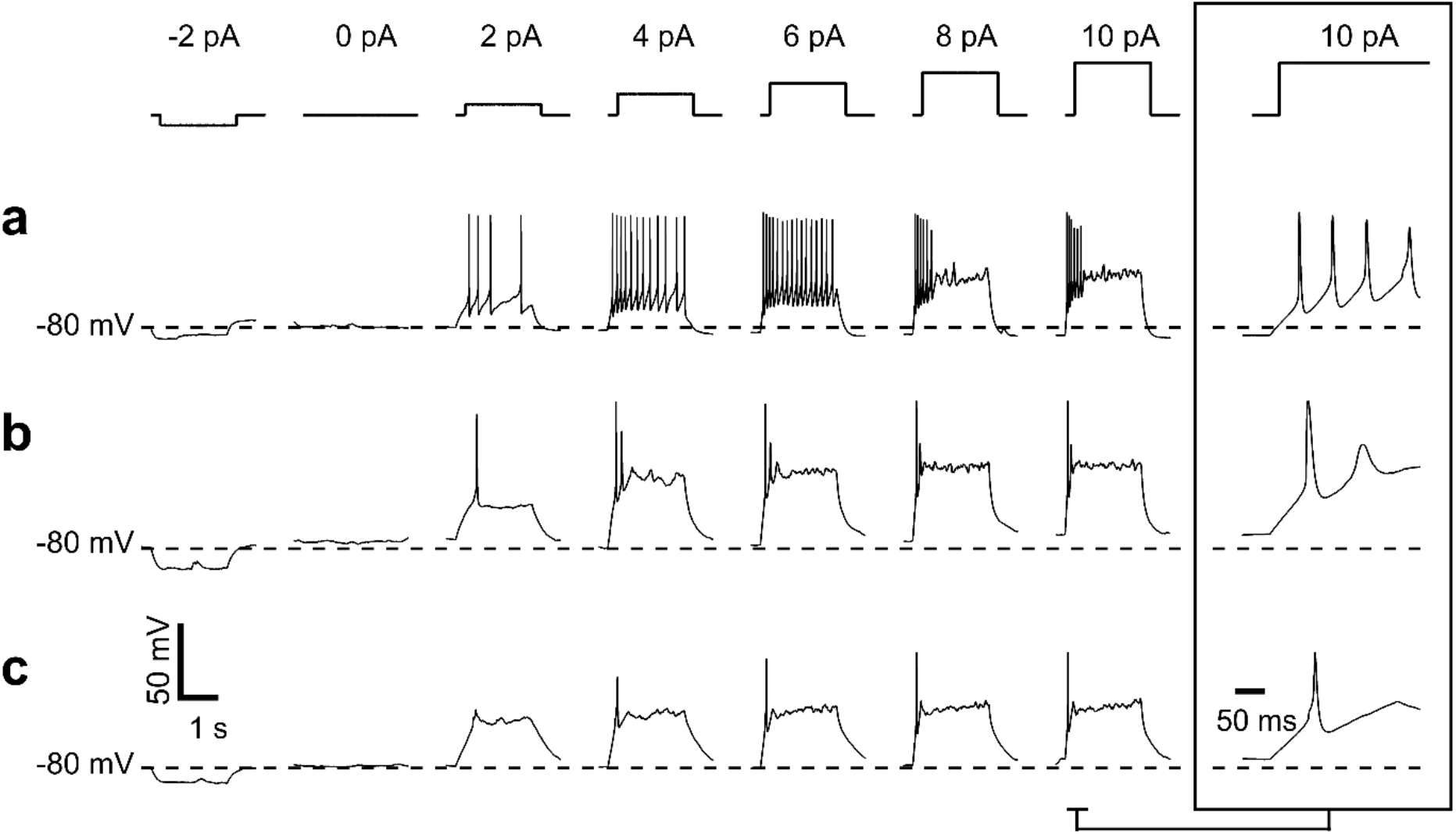
Firing patterns of olfactory sensory neurons recorded from acute slices of the human olfactory epithelium. **a-c** Spiking activity of three different olfactory sensory neurons recorded in whole-cell current-clamp in response to current steps of 2 s duration varying from -2 to 10 pA, with 2 pA increments, as indicated in the upper panel. Insets at the right show the details of firing activity generated with a 10 pA step and plotted on an expanded time scale for each cell.

These experiments show that whole-cell current-clamp experiments in slices from human olfactory epithelium are able to capture the electrophysiological heterogeneity in firing behavior of different olfactory sensory neurons, a characteristic common to other vertebrate species^35,62,63^.

### Responses to stimuli

To test if the olfactory transduction cascade is active in neurons of the human olfactory epithelium in acute slice preparation, we applied IBMX, a PDE inhibitor that acts on the transduction cascade by reducing the hydrolysis of cAMP. In the whole-cell voltage-clamp configuration at the holding potential of -80 mV, some neurons did not respond to stimulation with IBMX, although they generated an inward current when a solution containing high K^+^ was applied to test the neuron viability (Fig. 4a). Other neurons responded to 3 s stimulation of IBMX with an inward current that was slowly increasing its amplitude and then returning to baseline after IBMX removal (Fig. 4b). We found that 33 % (3 out of 9) of the neurons tested with IBMX displayed an inward current in response to IBMX with an average peak value of -37 ± 18 pA (n = 3), while the average value of the response to high K^+^ of the same neurons was -200 ± 17 pA (n = 3). The remaining 67% (6 out of 9) did not respond to IBMX although they responded to high K^+^.

**Fig. 4.**
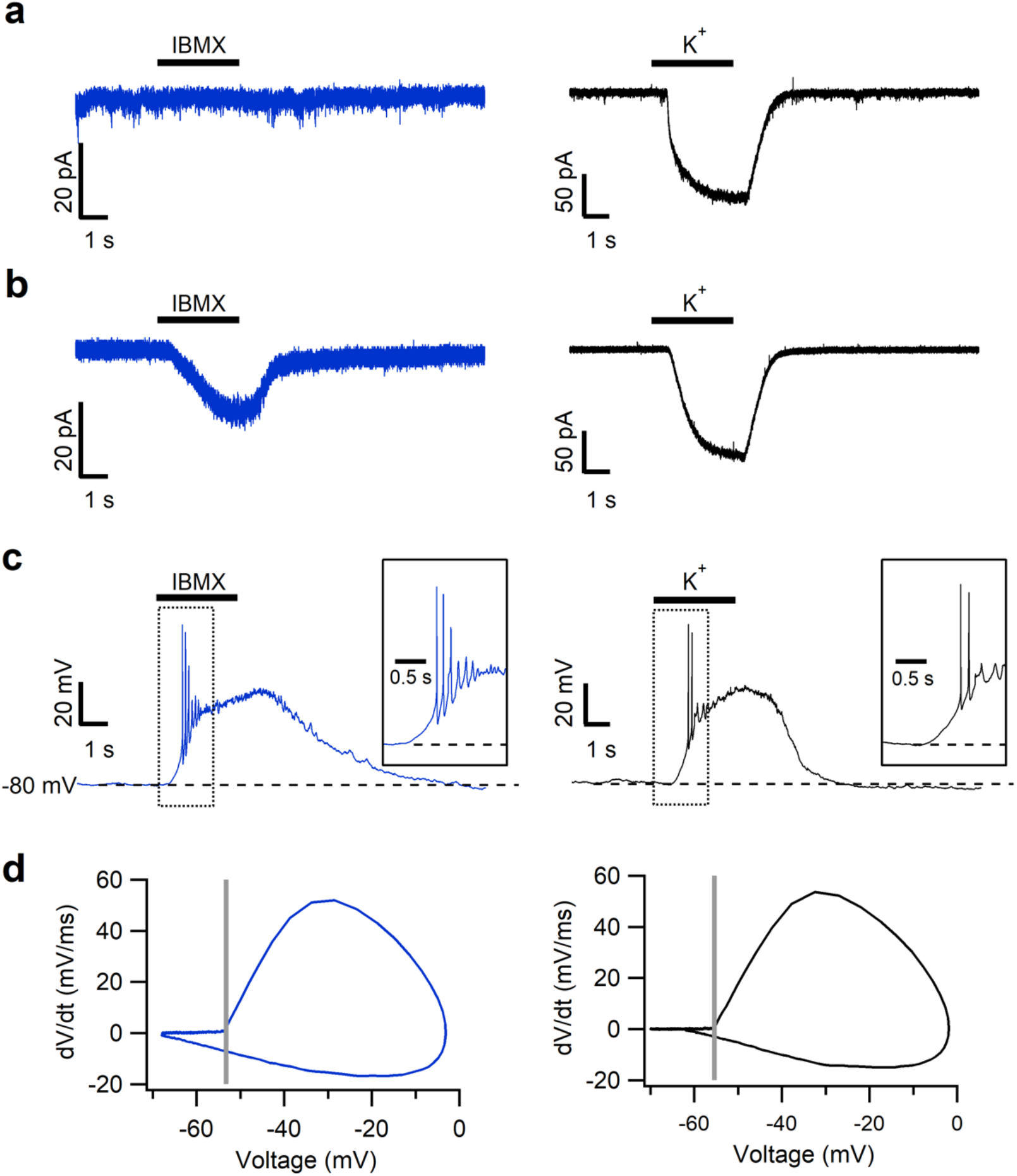
Responses of human olfactory sensory neurons to the phosphodiesterase inhibitor IBMX. **a** Example of an olfactory sensory neuron non-responding to 1 mM IBMX (blue trace; left) but responding to high K^+^ with an inward current (black trace; right). Holding potential was –80 mV. **b** Representative trace of an olfactory sensory neuron responding both to 1 mM IBMX and high K^+^ with an inward current. **c** Spiking activity measured in whole-cell current-clamp in the same olfactory sensory neuron shown in **b** stimulated with 1 mM IBMX or high K^+^. Insets show details of the spiking activity at the beginning of the stimulation on an expanded time scale. **d** Phase plots of the first action potentials from the responses shown in **c**. The crossing of vertical line with the upper loop indicates the voltage threshold for the first action potential. Stimulus duration for IBMX and high K^+^ was 3 s.

We also recorded the spiking pattern in response to IBMX or high K^+^ in the current-clamp configuration. The same neuron of Fig. 4b displayed firing both in response to IBMX and to high K^+^ (Fig. 4c). We analyzed the first action potential by using phase plot analysis, in which changes of membrane potentials with time (dV/dt) are plotted as a function of membrane potential (Fig. 4d). The action potential is represented by a loop, with the upper and lower parts representing the depolarization and repolarization phases, respectively. Phase plots were rather similar for the first action potential in both IBMX and high K^+^ with the following values (Fig. 4d): threshold -54 mV for IBMX and -55 mV for high K^+^; peak amplitude -3 mV for IBMX and -2 mV for high K^+^; the maximal rising slope of the depolarization phase was 52 mV/ms for IBMX and 54 mV/ms for high K^+^; the maximal rising slope of the repolarization phase was -17 mV/ms for IBMX and -15 mV/ms for high K^+^. Two other neurons responding to IBMX did not reach the voltage threshold for action potential generation.

To investigate if human olfactory sensory neurons in the slice preparation respond to odorants, we prepared two mixtures of odorants (mix 1 and mix 2, see Methods) and recorded current responses under the voltage-clamp configuration at a holding potential of -80 mV. We found that two out of five neurons that we considered viable, as they responded to high K^+^, responded to one of the two odorant mixtures with an inward transduction current. In one neuron, the peak amplitude of the current response was -18 pA with odorant mix 1 and -14 pA with IBMX, while mix 2 was not tested (Fig. 5a). In another neuron, odorant mix 1 did not activate any current, while mix 2 elicited an inward current of -27 pA peak amplitude (Fig. 5b).

**Fig. 5.**
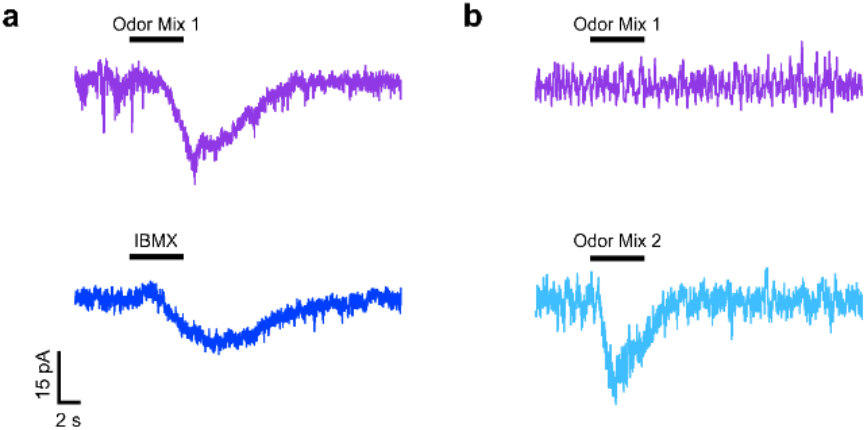
Responses of human olfactory sensory neurons to odorant mixtures. **a** One olfactory sensory neuron responding to odorant mix 1 and 1 mM IBMX with inward currents. **b** Another olfactory sensory neuron non-responding to odorant mix 1 but responding to odorant mix 2 with an inward current. Holding potential was – 80 mV and stimulus duration was 5 s.

Altogether, these results show that human olfactory sensory neurons in acute slices of the olfactory epithelium have an intact transduction cascade and can respond differently to odorant mixtures. The transduction cascade involves PDE and cAMP which acts as a second messenger, since application of IBMX produces inward currents and elicits action potentials.

## Discussion

In this study, we have provided the first demonstration that it is possible to obtain acute slices of the human olfactory epithelium from biopsies that are viable for electrophysiological experiments, crucial to unveil the molecular logic of the very first events of human olfaction.

The resting electrical properties, input resistance and resting membrane potential, that we measured from human olfactory sensory neurons in slices were found to be similar to those reported by Restrepo et al.^50^ from isolated human olfactory neurons and to those measured in other vertebrates^59^. However, in contrast to previous reports that recorded inward voltage-gated currents in only one out of eleven^50^ or one out of fourteen^58^ freshly dissociated human olfactory sensory neurons, we consistently recorded both voltage-gated transient inward currents and outward currents in olfactory neurons that were clearly visualized with a fluorescence dye. Our results suggest that avoiding enzymatic dissociation may be crucial for preserving the electrophysiological properties of these neurons. Therefore, recording from slices of the olfactory epithelium is a more suitable approach for studying the functionality of human olfactory sensory neurons.

In addition, we have also successfully recorded voltage-gated currents from human supporting cells and have found notable differences when compared with previous measurements taken in mice. First, recordings from supporting cells in mouse acute slices showed the presence of very large leak currents, which could be reduced by using gap junction blockers, while we did not observe any large leak currents in our measurements from human slices. Second, while mouse supporting cells displayed both voltage-gated transient inward currents and outward currents^60,61^, our measurements from human slices consistently showed only outward currents. Interestingly, TMEM16F is expressed in human supporting cells^15^, while in mice, it has only been found in the cilia of olfactory neurons^64^. These biophysical and molecular differences could have significant implications for our understanding of the contribution of supporting cells to the physiology in different species.

In human olfactory sensory neurons, we recorded action potential firing in the current-clamp configuration and observed that some human olfactory neurons responded to small depolarizing current steps of only 2-4 pA and 2 s duration with one or a few spikes, while others displayed a train of action potentials. Increasing current injections up to 10 pA some neurons displayed tonic firing in the range from 2 to 6 pA, while at higher current injections (8 and 10 pA) the same neurons showed brief action potential trains followed by a voltage plateau. Our findings are consistent with previous measurements in amphibians or rodents, indicating that different firing properties can be displayed also by human olfactory sensory neurons^35,62,63^.

Although the olfactory epithelium from amphibians and rodents has provided insights into the olfactory transduction, the mechanisms underlying this process in human olfactory sensory neurons still need to be understood. While transcriptomic data from the human olfactory epithelium have confirmed the expression of several genes known to be involved in the transduction cascade in rodent olfactory sensory neurons^8–11^, the expression and localization of the proteins have only been confirmed for the alpha and gamma subunits of the G protein using immunostaining data^11,13^. Here, we used immunohistochemistry and showed that AC3, the protein responsible for cAMP production, is expressed in the dendritic knob and cilia of human olfactory neurons, indicating a potential role of cAMP as a second messenger in olfactory transduction in humans, similar to what has been observed in rodents and amphibians. Furthermore, we demonstrated the localization of Ca^2+^-activated Cl^-^ channel TMEM16B in the dendritic knob and cilia, suggesting that it could have a significant role in human olfactory transduction, similar to previous findings in mice^34^.

By using patch-clamp recordings, we unveiled some crucial elements of the transduction in human olfactory sensory neurons. Specifically, we demonstrated that some olfactory neurons have a basal concentration of cAMP that is hydrolyzed by PDE in resting conditions. Indeed, when we applied the PDE inhibitor IBMX in whole-cell voltage-clamp, we recorded an inward current in some viable neurons, but not all. Not only we found that a cAMP increase induced by PDE inhibition with IBMX produced inward currents in voltage-clamp, but it also elicited action potential firing as measured in the current-clamp configuration. This indicates that cAMP build-up in olfactory sensory neurons can generate transduction currents capable of driving action potential firing.

Previous studies in mice have revealed that the constitutive activity of some odorant receptors leads to spontaneous transduction events in olfactory neurons, while other odorant receptors have a lower activity and do not induce spontaneous events^43–45^. Our experiments with IBMX suggest that human olfactory neuron may also exhibit spontaneous activity, depending on the specific odorant receptor expressed. In rodents, it has been shown that the spontaneous activity of odorant receptors is important to define the glomerular map in the olfactory bulb^65–67^. Whether this also occurs in human is an important question, although difficult to answer.

Recording odorant responses in olfactory sensory neurons is a challenging task, as each neuron only expresses one type of odorant receptor, which is activated by a limited number of odorants. To increase the probability of eliciting a response, we prepared two odorant mixtures that contained compounds known to be ethologically relevant for humans, including some that have previously been shown to activate human odorant receptors *in vitro*^68–70^. In whole-cell voltage-clamp, we recorded inward currents in response to each odorant mixture in different human neurons, thus showing that the transduction cascade initiated by odorant binding to specific receptor is fully functional in human olfactory sensory neurons in our slice preparation.

In summary, our data provide the first electrophysiological recordings of odorant responses in human olfactory sensory neurons from acute slices of the olfactory epithelium. We have demonstrated that the transduction mechanism involves PDE and that cAMP serves as a second messenger. Our findings lay the groundwork for future research using acute slices of the human olfactory epithelium, positioning humans as an ideal model for studying olfaction.

## Methods

### Human nasal tissue

Samples from human nasal tissue were obtained at the Section of Otolaryngology of the Department of Medical, Surgical and Health Sciences, University of Trieste, Trieste, Italy. The study was approved by the Ethics Committee on Clinical Investigation of the University of Trieste (nr 232/2016 and 110/2021), Friuli Venezia Giulia Region (CEUR-17236) and each patient provided written informed consent.

Biopsies were performed in the operating room from patients under general anesthesia at the end of the scheduled endoscopic sinonasal surgery, as previously reported^15^. In brief, two-three biopsy specimens were obtained from one nostril from the superior septum within the olfactory cleft and adjacent to the middle turbinate using a sickle knife and Blakesley forceps or cupped forceps. Once collected, biopsy specimens to be used for electrophysiology were immediately immersed in ice-cold artificial cerebrospinal fluid (ACSF) containing (in mM): 120 NaCl, 25 NaHCO_3_, 5 KCl, 1 CaCl_2_, 1 MgSO_4_, 10 HEPES, 10 glucose, pH 7.4, while those to be used for immunohistochemistry were immersed in paraformaldehyde (PFA) at 4% in PBS. Samples containing olfactory epithelium were obtained from 9 patients (5 males and 4 females, age between 24 and 70 years).

### Immunohistochemistry

As previously described^15^, human tissue samples used for immunohistochemistry were fixed in paraformaldehyde (PFA) at 4% in PBS pH 7.4 for 4 to 10 hours at 4 °C. After fixation, the tissue was kept in PBS pH 7.4 at 4 °C, typically from 2 to 24 hours. For cryoprotection of biopsies, the tissue was equilibrated overnight in 30% (w/v) sucrose in PBS at 4 °C. Then, the tissue was embedded in cryostat embedding medium (BioOptica) and immediately frozen at −80 °C. 16 μm sections were cut on a cryostat and mounted on Superfrost Plus Adhesion Microscope Slides (ThermoFisher Scientific). Sections were air-dried for 3 hours and used the same day or stored at -20 °C for later use. Cryostat embedding medium was removed from the tissue by incubating the slices in PBS for 15 minutes. The tissue was treated for 15 minutes with 0.5 % (w/v) sodium dodecyl sulfate (SDS) in PBS for antigen retrieval, then washed and incubated in blocking solution (5% normal donkey serum, 0.2% Triton X-100 in PBS) for 90 minutes and finally incubated overnight at 4 °C with the primary antibodies diluted in blocking solution. In the following day, the unbound primary antibodies were removed with PBS washes, then sections were incubated with Alexa Fluor conjugated secondary antibodies (1:500 dilution) in TPBS (0.2% Tween 20 in PBS) for 2 hours at room temperature, washed and mounted with Vectashield (Vector Laboratories) or FluoromontG (ThermoFisher). DAPI (5 mg/ml) was added to the solution containing secondary antibody to stain the nuclei.

The following primary antibodies (dilution; catalogue number, company) were used: polyclonal goat anti-OMP (1:1000; 019-22291, Wako), monoclonal mouse anti-β Tubulin III (TUJ1) (1:200; 801202, BioLegend), polyclonal rabbit anti-ERMN (1:200; NBP1-84802, Novus). polyclonal rabbit anti-AC3 (1:100; sc-588, Santa Cruz) and polyclonal rabbit anti-TMEM16B (1:200, NBP1-90739, Novus). The following secondary antibodies were used: donkey anti-rabbit Alexa Fluor Plus 594 (1:500; A32754, Life Technologies), donkey anti-rabbit Alexa Fluor 488 (1:500; A21206, Life Technologies), donkey anti-goat Alexa Fluor 647 (1:500; A32849, Life Technologies), donkey anti-mouse Alexa Fluor 594 (1:500, A-21203, Life Technologies), donkey anti-mouse Alexa Fluor 488 (1:500, A32766, Life Technologies).

Control experiments, excluding primary antibodies, were performed for each immunolocalization. We performed at least 2 independent human tissue replicates for each antibody tested. All attempts at replication were successful.

Z-stack images were acquired using NIS-Elements Nikon software at 1024 ×1024 pixels resolution of each single image and analyzed with ImageJ software (National Institute of Health, USA). Max projections of Z-stacks or individual images within the stacks were used to display results. Figure assembly was performed on ImageJ (National Institutes of Health) using ScientiFig plugin^71^. No image modification was performed other than brightness and contrast adjustment.

### Preparation of acute slices of human nasal tissue

Acute slices of human nasal epithelium used for electrophysiological experiments were prepared following a similar protocol to the one previously used for mouse olfactory and vomeronasal epithelium^15,61,72–76^. Within about 30 min from the biopsy, the human nasal epithelium was embedded in 3% Type I-A agarose (Sigma) prepared in ACSF once the agar had cooled to 38°C. Upon solidification, the agar block was fixed in a glass Petri dish and sliced with a vibratome (Vibratome 1000 Plus, Sectioning System) at 200 to 250 μm thickness in oxygenated ACSF solution. Slices were then left to recover for >30 min in chilled and oxygenated ACSF before electrophysiological experiments were initiated.

### Electrophysiological recordings

Slices were transferred to a recording chamber and continuously perfused with oxygenated (95% O_2_, and 5% CO_2_) ACSF by gravity flow. Each slice was anchored to the base of the recording chamber using a homemade U-shaped silver wire, holding down the agar support without touching the slice itself. Slices were viewed with an upright microscope (Olympus BX51WI) by infrared differential contrast optics with water immersion 20X or 60X objectives. The olfactory epithelium was easily distinguished from the respiratory one because the first had no moving cilia while the second had long beating cilia.

Whole-cell recordings were performed by patching the soma of the cells. Patch pipettes pulled from borosilicate capillaries (WPI) with a PC-10 puller (Narishige) had a resistance of 3–6 MΩ. The intracellular solution filling the patch pipette contained (in mM) 80 K-Gluconate, 60 KCl, 2 Mg-ATP, 10 HEPES, and 1 EGTA, adjusted to pH 7.2 with KOH. To visualize the morphology of the cell, 0.01mg/ml Alexa Fluor 594 carboxylic acid (Thermo Fisher, A33082) was dissolved in the patch pipette solution, diffused into the cell and the fluorescence image of the cell was observed under red fluorescence filter. Olfactory sensory neurons and supporting cells were clearly identified by their morphology (Fig. 2 a,d). Recordings in the whole-cell voltage-or current-clamp configurations were obtained with a Multiclamp 700B amplifier controlled by Clampex 10 via a Digidata 1440 (Molecular Devices). Data were low-pass filtered at 2 kHz and sampled at 10 kHz. Experiments were performed at room temperature (20–25°C).

Responses of olfactory sensory neurons to stimuli were tested with 1 mM 3-isobutyl-1-methylxanthine (IBMX) and two odorant mixtures. Mix 1 was composed of acetophenone, cineole, eugenol, heptaldehyde, isoamyl acetate, while mix 2 was composed of (R)-(-)-carvone, (S)-(+)-carvone, geraniol, hexanal, 7-hydroxycitronellal, (R)-(+)-limonene, octanal. Each odorant was present at 100 μM. For each experiment, the response to high K^+^ stimulation was used to evaluate the viability of the neuron and the time of stimulus application. Only olfactory sensory neurons that responded to high K^+^ solution were included in the analysis.

1 mM IBMX was prepared weekly by directly dissolving it in ACSF solution. For odorant mixtures, each odorant was dissolved in dimethyl sulfoxide (DMSO) to prepare stock solutions at 5 M and mixtures were prepared by diluting each odorant at a final concentration of 100 μM in ACSF on the day of the experiment.

Stimuli were focally delivered to the neuron through an 8-into-1 multibarrel perfusion pencil connected to a ValveLink8.2 pinch valve perfusion system (AutoMate Scientific). The tip of the perfusion head, with a diameter of 360 μm, was placed ∼500 μm away from the slice. To avoid mechanical artifacts, the slice was continuously perfused with ACSF and the flow out of the pipette was switched between ACSF and stimulus solutions.

All chemicals were purchased from Sigma-Aldrich unless otherwise stated.

### Data and statistical analysis

Igor Pro 8 software (WaveMetrics, Lake Oswego, OR, USA) was used for data analysis and figure preparation. All averaged data from individual experiments in different cells are presented as mean ± standard error of the mean (SEM) and number of cells (n). These data were normally distributed (Shapiro-Wilk test).

## Data availability

All relevant data are available from the corresponding authors.

## Author contributions

A.M., A.H.C and M.D. conceptualized the project and designed experiments. A.H.C. and K.G.V. performed immunohistochemistry and confocal microscopy. A.H.C., C.A.S.T., G.G., C.R., and F.A.M. made slices, performed patch-clamp recordings and data analysis. P.B.R., M.T., P.B. and G.T. collected human biopsies. A.M., M.D. and A.H.C. wrote the manuscript with comments from all the other authors.

## Statement of Ethics

Subjects have given their written informed consent and the study protocol has been approved by the Ethics Committee on Clinical Investigation of the University of Trieste (nr 232/2016, 110/2021) and Friuli Venezia Giulia Region (CEUR-17236).

